# Modeling acquired TKI resistance and effective combination therapeutic strategies in murine RET+ lung adenocarcinoma

**DOI:** 10.1101/2025.06.04.657911

**Authors:** Trista K Hinz, Anh T Le, Tristan Doan, Abby Ast, Sophia Jaramillo, Samantha Haines, Andre Navarro, Tejas Patil, Raphael A Nemenoff, Lynn E Heasley

**Author notes:** Corresponding Author: Lynn E. Heasley Department of Craniofacial Biology University of Colorado Anschutz Medical Campus Aurora, Colorado.

## Abstract

RET gene rearrangements yield oncogenic fusion proteins that drive a subset of lung adenocarcinomas (LUAD). The tyrosine kinase inhibitors (TKIs) selpercatinib and pralsetinib are approved therapies for RET+ lung cancers and have markedly improved clinical outcomes in these patients, but acquired resistance remains a hurdle to their durable management. Using a recently developed murine model of RET+ lung cancer driven by a *Trim24-Ret* fusion protein, two *Trim24-Ret* cell lines (TR.1 and TR.2) were established. Orthotopic lung tumors generated by transplantation of these cell lines initially respond to selpercatinib followed by prompt progression within ∼3 weeks of initiating TKI treatment. Cell lines derived from the selpercatinib-resistant TR.1 and TR.2 tumors exhibited *in vitro* sensitivity to MET and ERBB-targeted TKIs, indicating acquired bypass signaling through these receptor tyrosine kinases. Moreover, the selpercatinib-resistant TR.1 and TR.2 cell lines exhibited increased sensitivity to MEK and PTPN11 inhibitors relative to the parental cell lines, indicating a greater dependence on MAPK pathway signaling. The TKI-resistant cell lines showed no evidence for MET gene amplification, but exhibited varied induction of multiple genes that function within MET and ERBB2:ERBB4 interaction networks including ligands (HGF, NRG1), adaptors (GAB1) and co-receptors (NRP1). Consistent with an important role for MET signaling in driving acquired selpercatinib resistance, mice bearing orthotopic lung tumors derived from TR.1 or TR.2 cells that had progressed on selpercatinib treatment underwent significant re-shrinkage upon co-treatment with the MET inhibitor, crizotinib, although progression re-occurred. By contrast, upfront treatment with selpercatinib and crizotinib in orthotopic tumors yielded complete elimination of 78% of TR.1 tumors and a prolonged duration of response in TR.2 tumors. The findings highlight the failings inherent in treating acquired resistance mechanisms at progression and the potential therapeutic impact of predicting and targeting dominant mechanisms of resistance prior to or early after initiating oncogene-targeting TKI treatment in RTK-driven LUAD.

## Introduction

Lung adenocarcinoma (LUAD) presents with an array of oncogene drivers, many of which are gain-of-function mutants of members of the receptor tyrosine kinase (RTK) family of genes (1). In addition to oncogenic EGFR, ALK, ROS1 and NTRKs, gene rearrangements involving RET are detected in 1%-2% of LUAD tumors (2,3). Recently, the tyrosine kinase inhibitors (TKIs), selpercatinib (4) and pralsetinib (5) were developed and evaluated in RET+ lung cancer patients. The overall response rate in RET fusion-positive LUAD patients was greater than 70% in treatment-naïve patients with median progression-free survival approaching two years. Despite these impressive clinical responses, the general problem of acquired resistance remains a hurdle to long-term control or cure of LUAD driven by mutated RET in particular (6,7), and oncogenic RTKs in general (8,9).

Mechanisms of acquired TKI resistance include on-target missense kinase domain mutations that inhibit drug binding as well as selection for diverse bypass signaling pathways that circumvent dependency for the oncogenic RTK (7–9). While acquired resistance via on-target mutations is a general problem with TKIs deployed in RTK-driven lung cancers, selection for these mutations has been reduced with the advent of improved 2^nd^ and 3^rd^ generation agents designed to maintain potency on targets bearing such acquired resistance mutations. By contrast, drug resistance through emergence of bypass signaling pathways remains a hurdle to durable control of RTK-driven lung cancers despite improved TKIs. An extensive literature documents the activity of distinct RTKs including MET and ERBB family members to effectively engage in bypass signaling in the context of TKI therapy and is consistent with the nature of RTKs to function within complex co-activation networks (10). Bypass signaling can be achieved rapidly (minutes to hours) through wild-type EGFR in ALK, ROS1, RET and NTRK1-positive lung cancer cell lines (11). Also, the literature establishes MET as a common RTK mediating acquired resistance (12). At present, lung tumor progression following initial responses to oncogene-targeted agents is managed by serial monotherapies designed to target emergent resistance pathways. Clearly, this approach has not provided chronic control or cures. Notably, effective management of tuberculosis and HIV infections required up front combinations of agents, not serial monotherapies (13). In sum, despite a wealth of molecular information regarding mechanisms of acquired resistance to TKIs, clinical advance on how to design and deploy effective combination therapies in lung cancer remains difficult due to the many questions surrounding optimal drug combinations, when to deliver them in the course of therapy, and concerns regarding toxicity. It has been proposed that experimental models that permit discovery and prioritization of optimal drug combinations and the timing of their delivery are needed for advance on this general problem (14).

To investigate questions surrounding precision oncology in lung cancer, our group has developed murine lung cancer models and transplantable cell lines driven by oncogenic RTKs including EGFR, ALK and RET (15–17). When these murine lung cancer cell lines are used to establish orthotopic lung tumors in syngeneic hosts, TKI treatment yields a wide range in the depth and durability of therapeutic responses, similar to what is observed in TKI-treated LUAD patients. These responses are highly reproducible for individual cell lines, thus allowing preclinical studies to define mechanisms regulating depth of response and time to progression. Regarding RET+ lung cancer models, our recent study demonstrated modest shrinkage of orthotopic *Trim24-Ret* tumors in response to selpercatinib treatment promptly followed by acquired resistance and tumor progression (17). Thus, the *Trim24-Ret* cell line provides a new model to explore *in vivo* acquired resistance mechanisms and to test therapeutic strategies for improving or preventing resistance-mediated progression. The experiments presented in this study show that the signaling hubs comprised of HGF-MET and ERBB family members mediate *in vivo* acquired resistance to selpercatinib. Moreover, the studies demonstrate the superiority of up-front combinations of selpercatinib and the MET inhibitor, crizotinib relative to combining crizotinib at progression for obtaining durable tumor control. The results highlight the importance of identifying and targeting dominant resistance mechanisms prior to treatment or early in the course of TKI therapy rather than the failed strategy of intervening at the time of tumor progression.

## Materials and Methods

### Murine RET+ lung cancer cell lines

#### Murine Trim24-Ret lung cancer cell lines

The TR.1 cell line was generated from a *Trim24-Ret* tumor established in a male C57BL/6 Trp53^flox/flox^ mouse and has been previously described (17). Likewise, the previously described adenovirus encoding Cas9, Cre recombinase and the Trim24-Ret sgRNA pairs (17) were intratracheally instilled into a female C57BL/6 Trp53^flox/flox^ mouse. The resulting tumor was used to establish the cell line, TR.2. Both cell lines are TP53 null and are routinely propagated in RPMI-1640 medium supplemented with 5% fetal bovine serum, penicillin and streptomycin.

#### Establishment of cell lines from selpercatinib-resistant TR.1 and TR.2 orthotopic tumors

In **Supplementary Figure S1**, orthotopic TR.1 and TR.2 cell-derived lung tumors that had progressed on continuous selpercatinib treatment were excised, dissociated and propagated in full growth medium containing 200 nM selpercatinib to maintain the resistant phenotype. These cell lines are referred to as TR.1-1088, -1090, -1092, -1094 and TR.2-A0827, -A0828, -A0831 and -A0832.

#### Establishment of selpercatinib and pralsetinib-resistant TR.1 cell lines using in vitro methods

TR.1 cells were cultured in full growth medium containing 0.1% DMSO as a passage control or 30 nM selpercatinib or pralsetinib. Over the course of ∼60 days, the TKI concentrations were increased until cultures that readily grew in the presence of 400 to 500 nM TKI were obtained.

### *In vitro* cell growth and proliferation assays

#### 96-Well plate clonogenic assays

Suspensions (200 cells in 100 µl) of the cell lines were plated in 96-well tissue culture plates. The selpercatinib-resistant TR.1 and TR.2 cell lines were plated in medium containing 200 nM selpercatinib. Twenty-four hours later, 100 µl of medium containing 2X drug concentrations was added in triplicate to plates and incubated for 5-7 days. As a measure of cell number, DNA content was determined using the CyQUANT Direct Cell Proliferation Assay (Invitrogen, Carlsbad, CA) and a Biotek Synergy 2 fluorescence microplate reader (Agilent, Santa Clara, CA). Data are presented as percent of DMSO control-treated wells.

#### 6-well plate clonogenic growth assays

TR.1 and TR.2 cells were plated at 250 cells/well in 2 ml of growth medium in 6-well tissue culture plates. Twenty-four hours later, cells were treated with selpercatinib (0-300 nM) with or without 10 ng/ml recombinant murine HGF (Peprotech/Thermo Fisher Scientific, Waltham, MA). Fresh HGF was added every 3 days. Following 7-10 days of culture, the growth medium was aspirated, the wells were rinsed with phosphate-buffered saline and the cells were fixed and stained with 0.5 ml of 0.5% (w/v) crystal violet in 6.0% (v/v) glutaraldehyde for 30 minutes. The plates were rinsed extensively with distilled H_2_O, photographed and the total colony area per well was quantified with Metamorph imaging software (Molecular Devices, San Jose, CA). The data are presented as the percent of control DMSO-treated wells.

#### Population Doubling Assays

TR.1 and TR.2 cell lines were cultured in 10 cm plastic dishes in full medium supplemented with 0.1% DMSO as a control or selpercatinib (100 nM), afatinib (100 nM) or crizotinib (300 nM), alone and in combination. Every 3-4 days, the cells were trypsinized, counted and replated with fresh drugs. The population doubling level (PDL_t_) was calculated with the formula, PDL_t_ = 3.32 (log Cell Number Yield_t_ - log Cell Number Plated) + previous PDL.

### Immunoblotting analysis

Cells were collected in phosphate-buffered saline, centrifuged, and suspended in lysis buffer (0.5% Triton X-100, 50 mM β-glycerophosphate (pH 7.2), 0.1 mM Na_3_VO_4_, 2 mM MgCl_2_, 1 mM EGTA, 1 mM DTT, 0.3 M NaCl, 2 µg/ml leupeptin and 4 µg/ml aprotinin). Aliquots of the cell lysates containing 200 µg of protein were submitted to SDS-PAGE and immunoblotted with the following antibodies from Cell Signaling Technology (Danvers, MA): RET-pY905 (cs-3221), MET-pY1234/1235 (sc-3129), total MET (cs-3127), NRP-1 (cs-3725), pT202/pY204-ERK (cs-9101), pS473-AKT (cs-4060), total AKT (cs-9272), pY542-PTPN11 (cs-3751), pY580-PTPN11 (cs-3703), total PTPN11 (cs-3752), βActin (cs-4967). The total RET antibody (#134100) was purchased from Abcam (Cambridge, UK) and the total ERK antibody (sc-514302) was purchased from Santa Cruz Biotechnology (Dallas, TX).

### ELISA

Mouse HGF was measured in conditioned growth medium collected from selpercatinib-resistant TR.1 and TR.2 cell lines after 3 days of incubation with an ELISA kit purchased from R&D Systems (Minneapolis, MN). The levels are presented as pg HGF per µg of cellular protein.

### RNAseq

RNA was prepared from TR.1 cells cultured *in vitro* with DMSO or made resistant to selpercatinib or pralsetinib and submitted to the University of Colorado Cancer Center Genomics Shared Resource. Libraries were generated and sequenced on an Illumina NovaSeq 4000 to generate 2x151 reads. Fastq files were quality checked with FastQC, Illumina adapters trimmed with bbduk, and mapped to the mouse mm10 genome with STAR aligner. Counts were generated by STAR’s internal counter and reads were normalized to counts per million reads mapped (CPM) using the edgeR R package as previously described (18).

### Quantitative reverse transcription PCR

#### RNA

Cells were collected in 600 µL RNA Lysis Buffer and total RNA was purified using Quick-RNA MiniPrep kits (Zymo Research, Irvine, CA). Aliquots (5 µg) of total RNA were reverse transcribed in a volume of 20 µL using Maxima First Strand cDNA Synthesis Kit (Thermo Scientific, Pittsburgh, PA). Aliquots (2 µL) of a 1:5 dilution of the reverse transcription reactions were submitted to quantitative PCR in 10 µL reactions with SYBR Select Master Mix for CFX (Thermo Fisher Scientific) using a CFX Connect Real-Time PCR Detection System (BioRad, Hercules, CA). The following PCR primer pairs directed against the murine transcripts were used: MET (Fwd: 5’-GTTGTCCTTGGTGCAGAGGA-3’, Rev: 5’-CGTGAAGTTGGGGAGCTGAT-3’); HGF (Fwd, 5’-AGCTACAGAGGTCCCATGGA-3’, Rev, 5’-CAAGAACTTGTGCCGGTGTG-3’); NRG1 (Fwd: 5’-AGCTGGAGTAATGGGCACAC-3’, Rev: 5’-GCTGTGCCTGCTGTTCTCTA-3’); ERBB4 (Fwd: 5’-ACGAGCCTGCCCTAGTTCTA-3’, Rev: 5’-ATCGGTGCAAGGCTTACACA-3’); ERBB3 (Fwd: 5’-CTCGAGGAACACAGCATGGT-3’, Rev: 5’-CAGCCACACCAAAATCTGCC-3’); ERBB2 (Fwd: 5’-AGGCCCCAGGTGAATATCCT-3’, Rev: 5’-AAGACCAGTCTGGGTGCAAG-3’); EGFR (Fwd: 5’-ACCTCCATGCTTTCGAGAAC-3’, Rev: 5’-AGTGATGTGATGTTCAGGCC-3’) . Mouse GAPDH mRNA levels were measured (Fwd: 5’-CGTGGAGTCTACTGGCGTCTTCAC-3’, Rev: 5’-CGGAGATGATGACCCTTTTGGC-3’) as a housekeeper gene for normalization.

#### MET gene copy number

Genomic DNA was purified with Quick-DNA Miniprep kits (Zymo Research) according to the manufacturer’s protocol. Aliquots (2 µL) were submitted to quantitative PCR in 10 µL reactions with SYBR Select Master Mix and mouse MET primers (see above) or LINE-1 (Fwd: 5’-TAGCGAGTGACAGTTTGAGC-3’, Rev: 5’-TTCCGGTCCAGTGTTTCTTC-3’) as a reference gene.

### Orthotopic mouse lung tumor methods

C57BL/6J (#000664) mice were obtained from Jackson Laboratory (Bar Harbor, ME). All procedures and manipulations were performed under protocols approved by the Institutional Animal Care and Use Committees at the Rocky Mountain Regional VA Medical Center and University of Colorado Anschutz Medical Campus. TR.1 and TR.2 cells were injected into the left lobe of the lungs of mice. Cells were prepared in a solution of 1.35 mg/mL Matrigel (Corning #354234) diluted in Hank’s Balanced Salt Solution (Corning) for injection. Mice were anesthetized with isoflurane, the left side of the mouse was shaved, and a 1 mm incision was made to visualize the ribs and left lobe of the lung. Using a 30-gauge needle, 250,000 cells were injected in 40 µL of matrigel cell mix directly into the left lobe of the lung and the incision was closed with staples. Tumors were permitted to establish for 10 days and then the mice were submitted to micro-CT (µCT) imaging to obtain pre-treatment tumor volumes. Tumor-bearing mice were randomized into treatment groups (n = 10), either 20 mg/kg selpercatinib, 50 mg/kg crizotinib, the combination of the two drugs or diluent control (H_2_O) by oral gavage for 5 consecutive days followed by 2 days off. To monitor effects of drug treatment on tumor volume, mice were submitted to weekly µCT imaging by the University of Colorado Anschutz Medical Small-Animal IGRT Core using the Precision X-Ray X-Rad 225Cx Micro IGRT and SmART Systems (Precision X-Ray, Madison, CT). Tumor volume was quantified from µCT images using ITK-SNAP software36 (www.itksnap.org).

### Pharmacological agents

Pralesetinib, enbezotinib, capmatinib, sapitinib, afatinib, trametinib, gefitinib, AZD4547, pexidartinib, RMC-7977 and RMC-4550 were purchased from MedChemExpress (Monmouth Junction, NJ). Selpercatinib and crizotinib were generously made available by patient donation through the Thoracic Oncology unit, University of Colorado Hospital.

### Statistics

All graphing and statistical analyses were performed using GraphPad Prism version 10.4. Data are presented as the mean ± standard error of the mean (SEM).

### Data Availability

The RNAseq data generated in this study are deposited in GEO Datasets (GSE298990).

## Results

We recently described a strategy to generate murine lung cancers driven by *Trim24-Ret* and *Kif5b-Ret* fusion proteins catalyzed by CRISPR-Cas9-mediated gene translocations (17). In this study, a cell line was derived from a resulting *Trim24-Ret* lung tumor, TR.1, that yields orthotopic tumors following implantation into the left lung of syngeneic C57BL/6 mice. Orthotopic TR.1 tumors exhibited significant, but transient tumor shrinkage in response to LOXO-292/selpercatinib therapy. However, in contrast to murine EGFR and ALK-mutant LUAD models which exhibit durable responses to targeting TKIs (15,16), the TR.1 lung tumors exhibited progression on continued selpercatinib treatment within 3 weeks of initiating treatment. In the present study, we explored the mechanisms mediating rapid on-treatment progression of TR.1 tumors and tumors derived from a novel cell line, TR.2 and investigated strategies for extending durability of TKI responses.

### Derivation of selpercatinib-resistant cell lines from orthotopic tumors progressing *in vivo* on TKI

The selpercatinib treatment response of individual TR.1 and TR.2 orthotopic tumors as assessed by µCT is presented in **Supplementary Figure S1A and S1B**. Consistent with our recent study showing pronounced TKI responses compared to untreated controls (17), lung tumors established from both cell lines exhibited durable selpercatinib responses for ∼3 weeks followed by on-treatment progression. At the indicated time points, the progressing lung tumors were dissected and cell lines were established in growth medium containing 200 nM selpercatinib to maintain TKI-resistant phenotypes (see Materials and Methods). These cell lines are referred to as selpercatinib-resistant TR.1-1088, -1090, -1092, -1094 and TR.2-A0827, -A0828, -A0831, -A0832 cell lines throughout this study. In addition to selpercatinib, the TR.1 and TR.2 resistant cell lines exhibited resistance to the distinct RET inhibitors, pralsetinib and enbezotinib (**Fig. 1A and B**), a TKI that exerts activity against common RET resistance mutations (19). By contrast, parental TR.1 and TR.2 cells cultured in parallel in diluent DMSO maintained high sensitivity to all three RET inhibitors.

**Figure 1.**
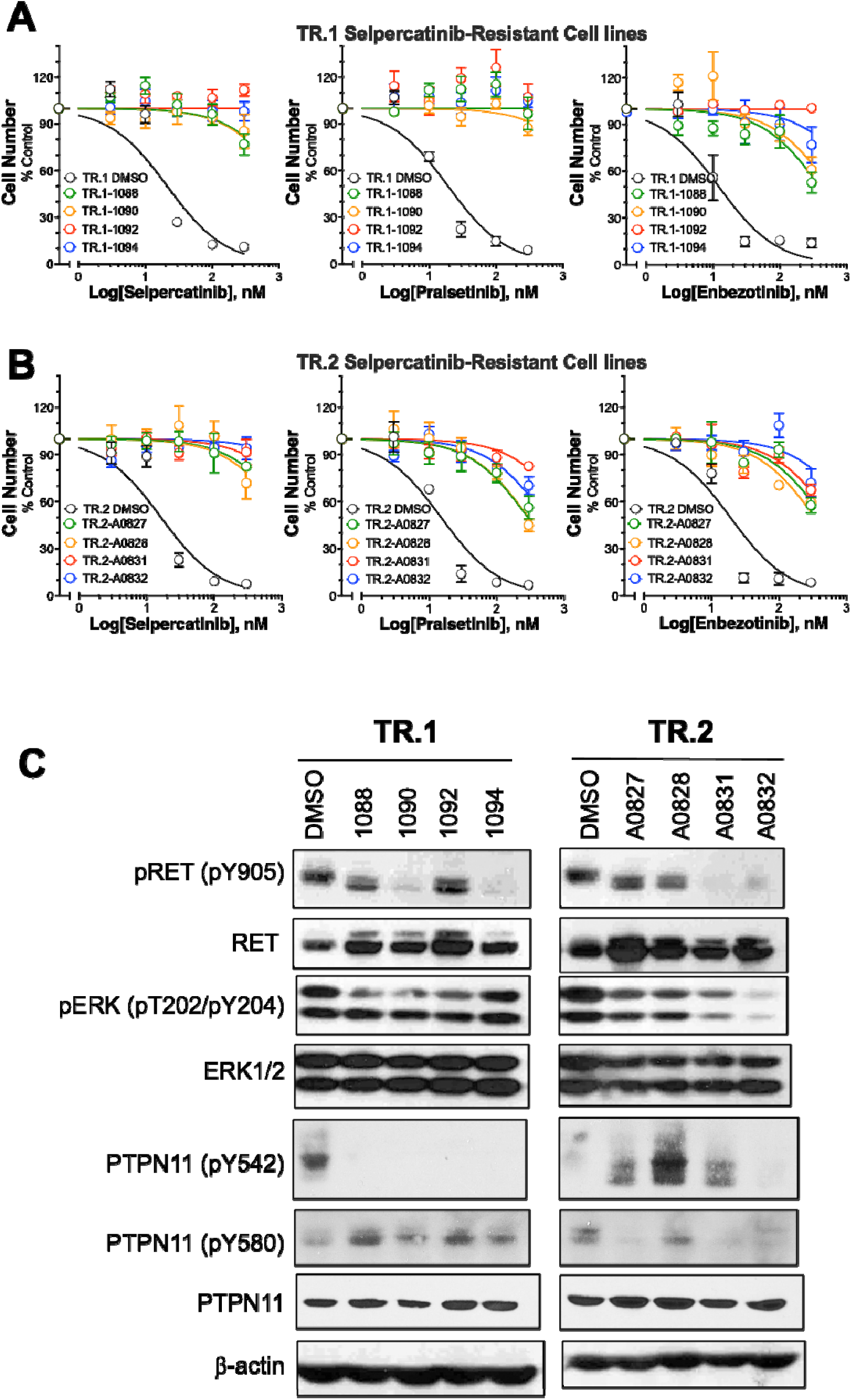
Cross-resistance of selpercatinib-resistant TR.1 and TR.2 cell lines to multiple RET-specific TKIs and variable retention of pY905-RET phosphorylation. Cell lines were derived from selpercatinib-resistant TR.1 (A) and TR.2 (B) tumors as described in the Materials and Methods. The cell lines were seeded in 100 µl at 200 cells/well of 96-well plates. Except for DMSO control TR.1 and TR.2 cells, all selpercatinib-resistant lines were seeded in medium containing 200 nM selpercatinib. The next day, 100 µl of 2X concentrated drug stocks in full medium was added in triplicate and the plates were incubated for 5-7 days. Relative cell number was assayed with CyQUANT reagent as described in the Materials and Methods. The data were submitted to curve fitting (log inhibitor vs. normalized response and robust regression) using Prism 10 and the IC_50_ values are presented in Supplementary Table S1. The data are the mean and SEM of 3 or more independent experiments. C, Cell-free extracts were prepared from the indicated control and selpercatinib-resistant cell lines (cultured in 200 nM selpercatinib) and submitted to SDS-PAGE. Following electrophoretic transfer, the filters were immunoblotted with antibodies to pY905-RET, total RET, pERK, total ERK, pY542- and pY580-PTPN11, total PTPN11 and β-actin as a loading control.

Immunoblot analysis of the selpercatinib-resistant cell lines cultured continuously in 200 nM selpercatinib demonstrated that *Trim24-Ret* protein was expressed at levels equal to that observed in parental TR.1 and TR.2 cells (**Fig. 1C**). Phospho-Y905 RET levels were markedly reduced compared to parental cells in some of the resistant cell lines (TR.1-1090 and -1094, TR.2-A0831 and -A0832), but not TR.1-1088 and -1092 or TR.2-A0827 and -A0828. While inhibition of Y905 phosphorylation located in the activation loop of TRIM24-RET is anticipated due to the inclusion of selpercatinib to the growth medium, retention of pY905-RET could indicate the acquisition of secondary missense mutations that prevent TKI binding. However, direct sequencing of the PCR-amplified murine *Trim24-Ret* kinase domain sequences detected no missense mutations that might confer TKI resistance (data not shown). The findings reveal that the selpercatinib-resistant cell lines are also insensitive to two distinct RET-specific TKIs, pralsetinib and enbezotinib, and retain expression of the *Trim24-Ret* oncogene.

### TKI and RAS-MAPK pathway inhibitor screen in selpercatinib-resistant TR.1 and TR.2 cell lines

Initial characterization of the selpercatinib-resistant TR.1 and TR.2 cell lines (**Fig. 1 and Suppl. Fig. S1**) supports bypass signaling as a likely mechanism of TKI resistance since no on-target mutations were identified. To functionally screen for dominant bypass signaling pathways, the selpercatinib-resistant TR.1 and TR.2 cell lines were tested for sensitivity to multiple TKIs with specificity for distinct RTKs. Clonogenic growth assays were performed in a 96-well format with a range of drug concentrations permitting the calculation of IC_50_ values which are summarized in **Supplementary Table S1**. The TR.1 and TR.2 selpercatinib-resistant cell lines exhibited acquired sensitivity to MET-targeting TKIs, crizotinib and capmatinib relative to the parental cell lines (**Figure 2A and B**). Notably, the MET TKIs exerted only partial reduction in cell growth with the magnitude of the inhibitory response being greater in the TR.1 selpercatinib-resistant cell lines (∼30 to 60% growth reduction) compared to the TR.2 cell lines. The selpercatinib-resistant cell lines also exhibited partial sensitivity to two distinct pan-ERBB inhibitors, sapitinib and afatinib (**Fig. 2A and B**), but not the EGFR-specific inhibitor, gefitinib (**Suppl. Table S1**), indicating a role for ERBB2, ERBB3 and/or ERBB4 in selpercatinib resistance. Variation amongst the different selpercatinib-resistant cell lines was observed. For example, TR.1-1092 cells exhibited little or no sensitivity to sapitinib and afatinib and were the most sensitive to the MET TKIs (**Fig. 2A**). By contrast, TR.1-1088 and -1090 were the least sensitive to the MET inhibitors and the most sensitive to ERBB inhibitors amongst the TR.1 resistant lines. The TR.2 selpercatinib-resistant lines exhibited a greater magnitude of growth inhibition by the ERBB pathway inhibitors than the TR.1 lines (**Fig. 2B**). Finally, FGFR and CSF1R-specific inhibitors AZD4547 and pexidartinib, respectively, exhibited no growth inhibition in parental or resistant cell lines (**Suppl. Table S1**).

**Figure 2.**
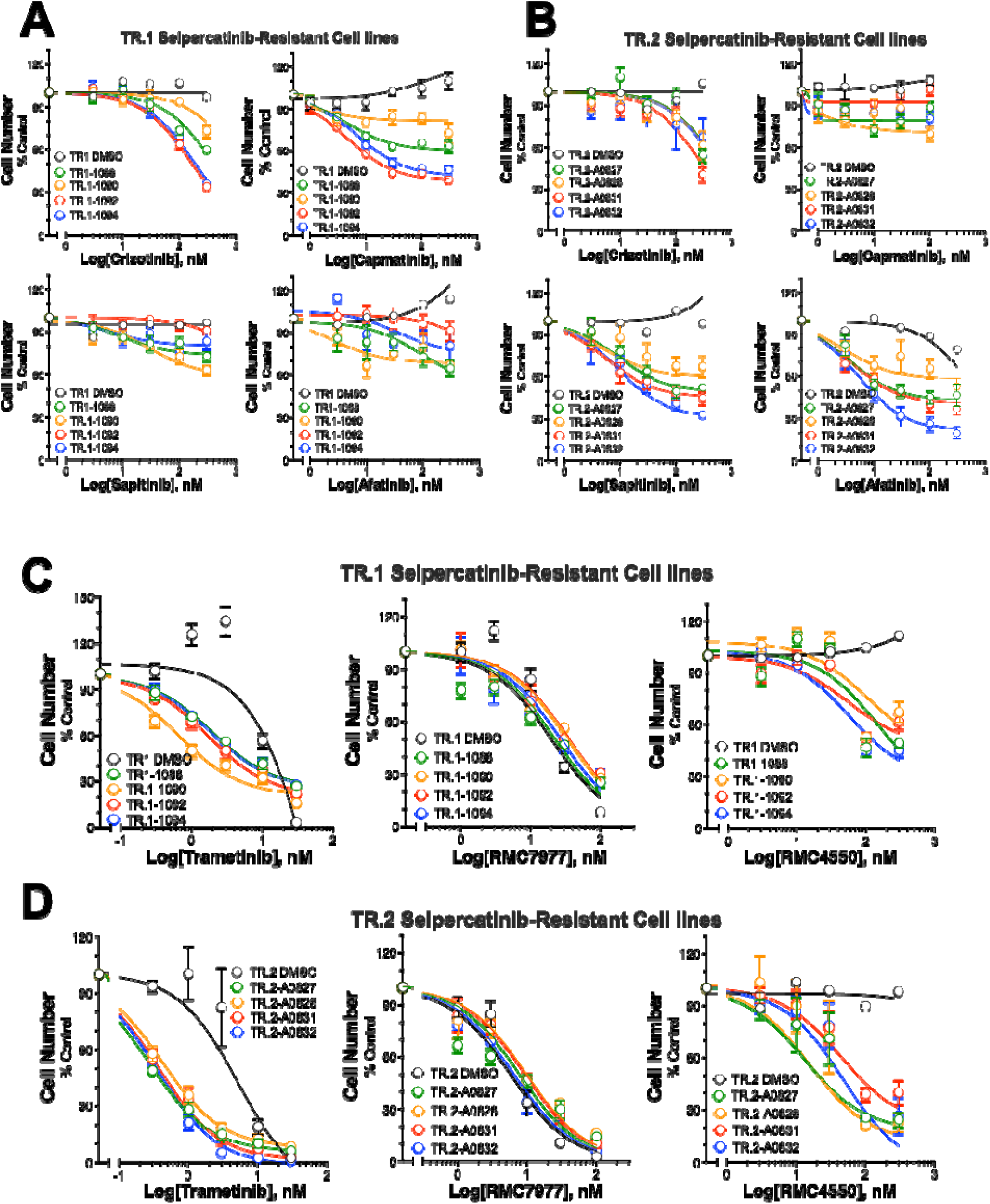
Pharmacological inhibitor screen of selpercatinib-resistant TR.1 and TR.2 cell lines. Control TR.1 and TR.2 cells cultured in DMSO and the cell lines derived from selpercatinib-resistant tumors (cultured in 200 nM selpercatinib) were seeded in 96-well plates at 200 cells/well and incubated for 5-7 days with MET inhibitors (crizotinib, capmatinib), ERBB inhibitors (sapitinib, afatinib), MEK inhibitor (trametinib), pan-RAS inhibitor (RMC7977) or PTPN11/SHP2 inhibitor (RMC4550) at the indicated doses. Relative cell numbers were measured with CyQUANT reagent (see Materials and Methods) and the data were analyzed with Prism 10. The IC_50_ values are presented in Supplementary Table S1.

The selpercatinib-resistant cell lines were also tested for MAPK pathway dependency based on sensitivity to the MEK inhibitor, trametinib, the pan-RAS inhibitor, RMC-7977 (20), and the PTPN11/SHP2 inhibitor, RMC-4550 (21). Parental TR.1 and TR.2 and the selpercatinib-resistant derivative cell lines exhibited equivalent sensitivity to pan-RAS inhibitor RMC-7977, but the selpercatinib-resistant TR.1 and TR.2 cell lines exhibited ∼15-30-fold enhanced sensitivity to trametinib (**Fig. 2C** and **D, Suppl. Table S1**). Increased trametinib sensitivity has been previously observed in RET-dependent LC-2/ad lung cancer cells rendered resistant to ponatinib, a multi-kinase TKI with affinity for RET (22). Moreover, the selpercatinib-resistant lines were uniformly sensitive to the PTPN11 inhibitor RMC-4550 while parental TR.1 and TR.2 cells were fully insensitive over the range of concentrations tested (**Fig. 2D**), demonstrating an acquired vulnerability to this agent in the context of selpercatinib-resistance. The molecular basis of this enhanced sensitivity to trametinib and RMC-4550 is not associated with increased pERK levels in the selpercatinib-resistant cell lines (**Fig. 1C**). While total PTPN11 levels were not different between control and selpercatinib-resistant TR.1 and TR.2 cells (**Fig. 1C**), phosphorylation of two C-terminal PTPN11 tyrosine residues, Y542 and Y580 with putatively positive effects on enzyme activation and adaptor protein recruitment (21) exhibited rather varied alterations assessed on immunoblots (**Fig. 1C**). PTPN11-pY542 was undetectable in the selpercatinib-resistant TR.1 cell lines compared to TR.1 control cells, but clearly detectable in 3 of the 4 selpercatinib-resistant TR.2 cell lines. By contrast, PTPN11-pY580 was modestly increased in the selpercatinib-resistant TR.1 cell lines, but reduced in the selpercatinib-resistant TR.2 cell lines. The findings in **Figure 2C** and **D** support increased reliance on PTPN11 and ERK signaling, but not RAS function for growth and survival in the setting of selpercatinib resistance, although this is not explained by increased pERK or pY-PTPN11 status.

To determine if the acquired resistance to selpercatinib is reversible, TR.1-1088 and -1092 selpercatinib-resistant cell lines were cultured in the absence of selpercatinib for 14 days. As shown, these cell lines became partially, but not fully re-sensitized to selpercatinib (**Suppl. Fig. S2**). Moreover, the increased sensitivity to crizotinib, afatinib and the MAPK pathway inhibitors, trametinib and RMC-4550 was fully reversed. The reversibility of the TKI-resistant phenotype is consistent with transcriptional induction of one or more bypass signaling pathways.

### Mechanisms mediating acquired sensitivity to MET and ERBB pathway inhibitors in selpercatinib-resistant cell lines

During the generation and evaluation of the selpercatinib-resistant cell lines derived from orthotopic tumors, selpercatinib and pralsetinib-resistant TR.1 cells were generated through standard *in vitro* culture methods involving stepwise increases in TKI concentrations. Similar to the TR.1 selpercatinib resistant cell lines derived from tumors, *in vitro* resistant cell lines able to grow in ∼500 nM TKI exhibited evidence for acquired sensitivity to crizotinib and afatinib (**Suppl. Fig. S3A).** To explore potential resistance mechanisms, total RNA was prepared from replicate selpercatinib and pralsetinib-resistant *in vitro* cultures as well as DMSO-treated TR.1 cells as a control. RNA was submitted to bulk RNAseq (see Materials and Methods) and expression of candidate RTK bypass pathways was interrogated, revealing evidence for increased expression of multiple genes that are predicted to function within the MET and ERBB2 interaction networks defined by the STRING database (string-db.org/; **Suppl. Fig. S3B and C**). Among the top 10 MET interactors, mRNA expression levels of MET, HGF, GAB1, PLXNB1 and LRIG1 were increased relative to the DMSO control cells (**Suppl. Fig. S3D).** Also, the HGF and PLXNB1 interactor, NRP1 (23,24) was induced in the selpercatinib and pralsetinib-resistant TR.1 lines derived *in vitro*, but ERBB3 expression was markedly diminished. While important for propagation of MET signaling, expression levels of CBL, SRC, HRAS and GRB2 were not changed in the TKI-resistant cells. The increased HGF expression predicted by the RNAseq data was validated by ELISA of conditioned medium (**Suppl. Fig. S3E**). Regarding the ERBB2 pathway, the RNAseq data revealed upregulation of ERBB4 and the EGF family ligand, neuregulin 1 (NRG1) mRNA levels in the TKI-resistant TR.1 cell lines (**Suppl. Fig. S3C and D**). Notably, NRG1 is a ligand for ERBB2:ERBB3 and ERBB2:ERBB4 heterodimers and ERBB4:ERBB4 homodimers (25). By contrast, expression levels of the majority of EGF family ligands were decreased in the *in vitro* TKI-resistant TR.1 cells (data not shown). Among the top 10 ERBB2 interactors (**Suppl. Fig. S3C**), EGFR and ERBB2 expression were unchanged, but ERBB3 and ERRFI1, a negative regulator of EGFR and ERBB2 (26,27), were markedly downregulated (**Suppl. Fig. S3D**). Expression of the signaling adaptors GRB2 and SHC1 and the FAK family member PTK2 were not changed, but PIK3R1 was increased in the TKI-resistant TR.1 cells. These findings support the hypothesis that *in vitro* acquisition of selpercatinib and pralsetinib resistance in TR.1 cells is mediated by increased expression of multiple genes that function within MET and ERBB2 interaction networks rather than through increased expression of a single dominant driver gene.

Based on these data, the expression levels of MET, HGF and selected interacting proteins were explored in the selpercatinib-resistant TR.1 and TR.2 cell lines derived from orthotopic tumors. Modestly elevated MET mRNA levels were noted in the selpercatinib-resistant TR.1 cell lines, but not the TR.2 lines (**Fig. 3A**). Increased HGF mRNA (**Fig. 3A**) and protein (**Fig. 3C**) were observed in TR.1-1092 and -1094, the most sensitive lines to crizotinib and capmatinib (**Fig. 2A**), but not in the TR.2 selpercatinib-resistant lines. No changes in MET gene copy number assessed by qPCR analysis of genomic DNA (**Fig. 3B**) or MET protein measured by immunoblotting were observed (**Fig. 3D**) in the TR.1 and TR.2 selpercatinib-resistant cell lines relative to control cells. All four selpercatinib-resistant TR.1 cell lines exhibited increased pY1234/1235-MET relative to control TR.1 cells with extracts from TR.1 cells treated for 2 hours with exogenous mouse HGF as a positive control (**Fig. 3D**). By comparison, pY1234/1235-MET in the selpercatinib-resistant TR.2 cells was detectable, but similar to that observed in untreated control TR.2 cells. In addition, protein expression of the HGF interactor NRP1 (23,24), was increased in all four selpercatinib-resistant TR.1 cell lines and also in TR.2-A0827 and -A0828 cells (**Fig. 3D**). To test if exogenous HGF can drive selpercatinib resistance, parental TR.1 and TR.2 cells were treated with a range of selpercatinib doses with and without recombinant HGF (10 ng/ml) in a clonogenic growth assay (see Materials and Methods). As shown in **Figure 3E**, exogenous HGF fully prevented growth inhibition by selpercatinib in TR.1 cells and significantly reduced TKI sensitivity in TR.2 cells, indicating that either autocrine or paracrine HGF-induced signaling through MET is capable of serving as a bypass pathway in these cell lines.

**Figure 3.**
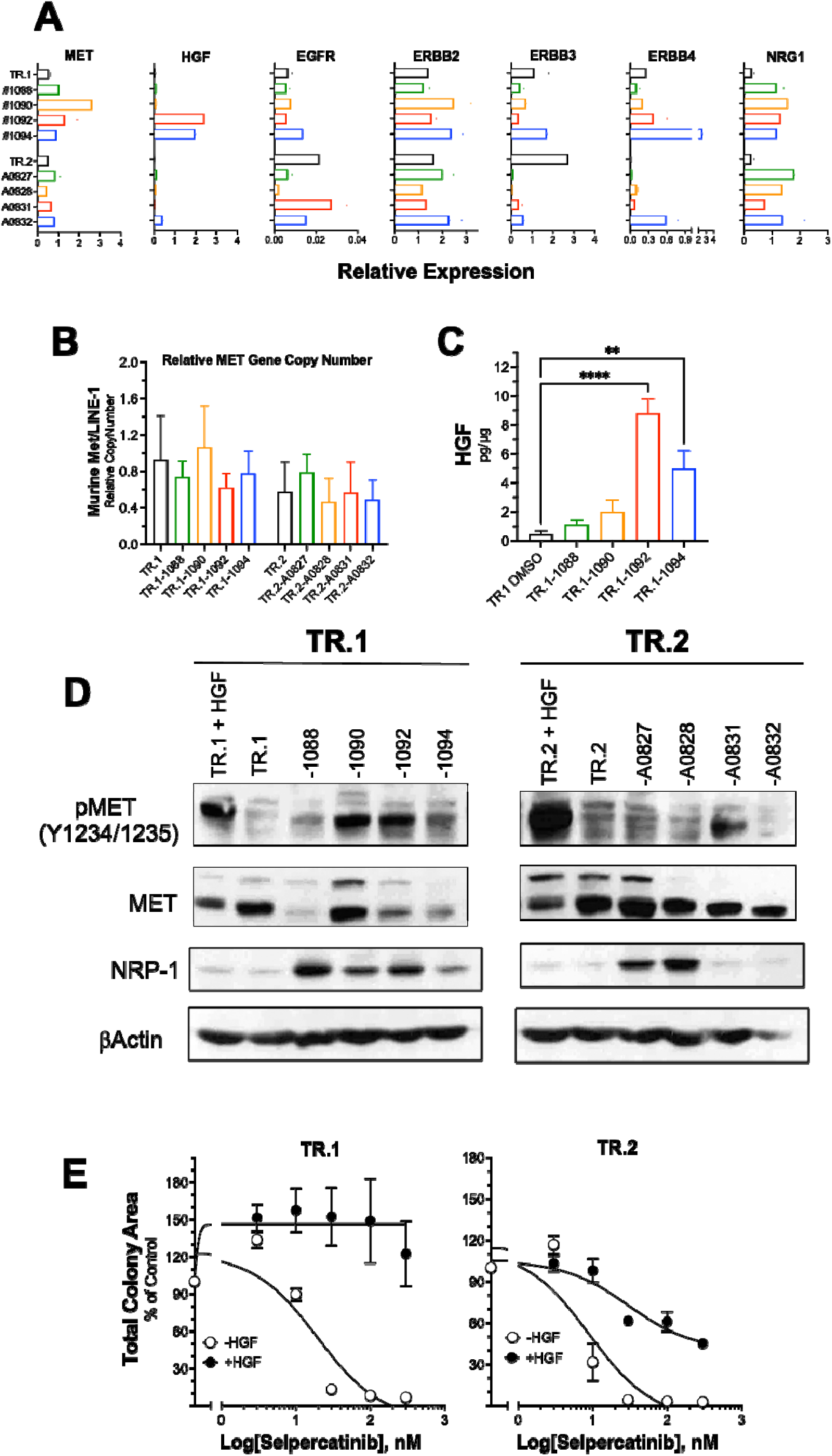
Induction of components of MET and ERBB interaction networks in selpercatinib-resistant TR.1 and TR.2 cells. **A**, RNA was purified from the selpercatinib-resistant and parental TR.1 and TR.2 cell lines and following reverse-transcription, submitted to qPCR with primer pairs specific to murine MET, HGF, EGFR, ERBB2, ERBB3, ERBB4 and NRG1. The data were normalized to Z-scores and are presented in a heatmap format. **B**, Genomic DNA was prepared from the indicated cell lines and submitted to qPCR analysis for murine MET and LINE-1 as an internal control. The MET gene copy number values were normalized to the LINE-1 levels in the samples and are presented as the mean and SEM of three independent experiments. **C**, Conditioned medium from the selpercatinib-resistant TR.1 cells was submitted to ELISA for HGF. The values are normalized to total cellular protein and are the means and SEM of 3 independent experiments. **D**, Cell-free extracts from TR.1 and TR.2 cells treated for 2 hours with 10 ng/ml HGF and the selpercatinib-resistant cell lines were submitted to SDS-PAGE and immunoblotted with antibodies to the indicated proteins. **E**, Parental TR.1 and TR.2 cells were seeded in 6-well plates and cultured with increasing doses of selpercatinib in the presence or absence of 10 ng/ml HGF. After 7-10 days, the resulting colonies were stained with crystal violet and quantified as described in the Materials and Methods.

In addition to evidence for increased MET pathway activity, qPCR verified increased mRNA expression of the ERBB ligand, NRG1, in all of the selpercatinib-resistant TR.1 and TR.2 cell lines relative to parental controls (**Fig. 3A**). Similar to the *in vitro* selpercatinib/pralsetinib-resistant TR.1 cells (**Suppl. Fig. S3**), expression levels of EGFR and ERBB2 mRNAs were not increased in the selpercatinib-resistant TR.1 and TR.2 cell lines. In fact, EGFR mRNA levels were markedly lower in TR.2-A0827 and -A0828 cells which exhibited the lowest sensitivity to sapitinib and afatinib (**Fig. 2B**). ERBB3 mRNA was generally decreased in the selpercatinib-resistant TR.1 and TR.2 cell lines and ERBB4 mRNA was increased in TR.1-1094 and TR.2-A0832 cells (**Fig. 3A**) where TR.1-A0832 cells are the most sensitive to ERBB-specific TKIs (**Fig. 2B**). The data support the hypothesis that in vivo acquired resistance of TR.1 and TR.2 tumors to selpercatinib is mediated by induction of multiple genes whose products function within the MET and ERBB2 interaction networks.

### Testing of TKI combinations to prevent bypass pathway-mediated resistance

#### In vitro population doubling experiment

To test the role of HGF-MET and ERBB pathways in the acquisition of selpercatinib resistance modeled *in vitro*, TR.1 and TR.2 cells were cultured with 100 nM selpercatinib in the presence and absence of 100 nM afatinib or 300 nM crizotinib. Treatment with diluent DMSO, afatinib or crizotinib, alone, was performed to test single agent activity of the ERBB and MET inhibitors. Every 3 to 4 days, the cell cultures were trypsinized, counted and replated to determine the population doubling rate (see Materials and Methods). Relative to DMSO or single-agent afatinib or crizotinib, 100 nM selpercatinib induced a decrease in overall cell number in both lines (see day 8 in **Fig. 4**). Combining afatinib with selpercatinib yielded a greater decrement in cell number than selpercatinib, alone, especially in TR.2 cells at days 8-14. By contrast, combining crizotinib with selpercatinib did not inhibit cell proliferation more than selpercatinib alone within the first week of the experiment. By day 14-18 of the experiment, the population doubling of cells treated with selpercatinib alone or in combination with afatinib recovered and achieved a rate similar to that observed in DMSO-treated cells (**Fig. 4**). By contrast, TR.1 and TR.2 cells treated with combined selpercatinib and crizotinib exhibited reduced population doubling rates from day 14 till the end of the experiment despite the lack of initial combination effects at day 8 of the experiment. The results demonstrate that combining afatinib with selpercatinib yields more marked early growth inhibition, but acquired resistance occurs promptly. By contrast, combining selpercatinib and crizotinib yielded no immediate effect on growth inhibition beyond that provided by selpercatinib (before and up to day 8), but did provide prolonged suppression of population doubling in both TR.1 and TR.2 cells relative to selpercatinib alone.

**Figure 4.**
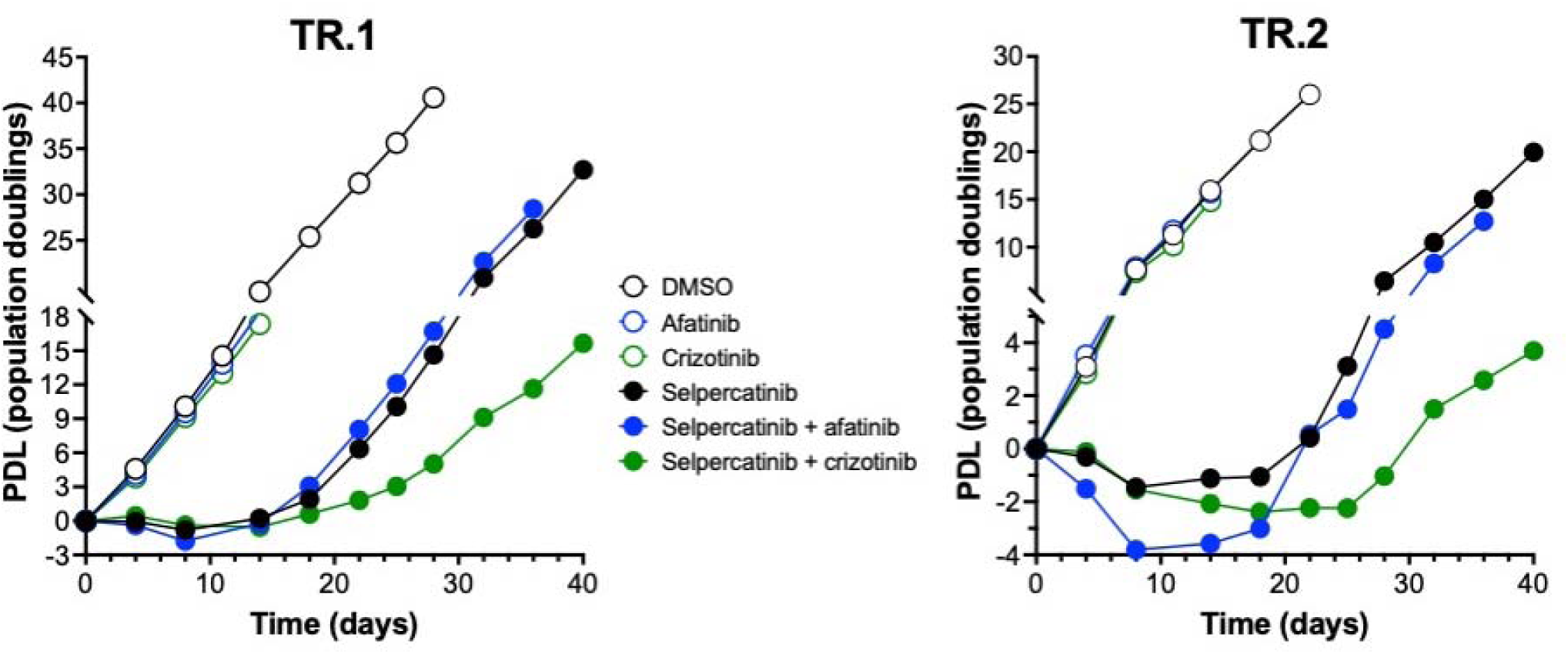
Time course of *in vitro* single agent and combination TKI responses in TR.1 and TR.2 cells. TR.1 and TR.2 cells were plated at defined cell numbers in 10 cm dishes and treated with diluent DMSO as a control, afatinib (100 nM), crizotinib (300 nM), selpercatinib (100 nM), selpercatinib plus afatinib or selpercatinib plus crizotinib. The cultures were trypsinized, counted and replated every 3-4 days and the population doubling values were calculated as described in the Materials and Methods. The experiment shown is representative of another independent experiment.

To explore early signaling effects of the TKIs, TR.1 and TR.2 cells were treated for 2 to 24 hrs with selpercatinib alone and in combination with afatinib or crizotinib (**Suppl. Fig. S4A and B**). Immunoblot analysis revealed that the dose of selpercatinib (100 nM) used in the population doubling experiment was sufficient to completely block pY905-RET phosphorylation, but residual pERK was clearly detectable in TR.1 and TR.2 cells and both pERK and pS473-AKT levels increased at the 24-hour timepoint, suggesting rapid ERK and AKT pathway reactivation. Combining selpercatinib and afatinib yielded greater pERK and pAKT inhibition relative to single agent selpercatinib in both TR.1 and TR.2 cells. At these early time points, combined selpercatinib and crizotinib treatment failed to reduce pERK and pAKT levels more than selpercatinib alone. The enhanced early growth inhibition and stronger reduction of ERK and AKT pathway activity with combined selpercatinib and afatinib is consistent with findings that the EGFR/ERBB pathway can function as an immediate bypass mechanism that reduces growth inhibition through multiple RTK fusions in lung cancer cells (11). The lack of effect of combined crizotinib and selpercatinib at early time points is consistent with the population doubling experiment in **Figure 4**.

#### In vivo activity of selpercatinib and crizotinib combinations

The results in **Figure 4** support the combination of crizotinib and selpercatinib as a superior therapy relative to selpercatinib and afatinib for treating orthotopic *Trim24-Ret*-driven lung tumor models. To advance these findings into the *in vivo* setting, mice bearing orthotopic TR.1 or TR.2 lung tumors were treated with selpercatinib alone or in combination with crizotinib as described in the Materials and Methods. Treatment with single agent selpercatinib induced significant tumor shrinkage to ∼25% of the initial volume followed by progression after day 20 in TR.1 tumors (**Fig. 5A**). Single agent selpercatinib induced TR.2 tumor shrinkage to ∼50% of initial volume and tumor progression was clearly evident by day 26 of treatment (**Fig. 5C**). At this time, addition of crizotinib (50 mg/kg) with maintenance of selpercatinib induced re-shrinkage of the progressing TR.1 and TR.2 tumors, although regrowth promptly occurred with both tumor models (**Fig. 5A-D**). Relative to single agent selpercatinib, up-front treatment with selpercatinib and crizotinib induced deeper tumor shrinkage in both the TR.1 and TR.2 tumors (**Fig. 5A and C**). Moreover, there was no evidence of acquired resistance or progression in TR.1 tumors (**Fig. 5A**), although TR.2 tumors treated with up-front selpercatinib and crizotinib began to re-grow after 29 days of therapy (**Fig. 5C and D**). After 48 days of up-front combination therapy, 7 of the 9 TR.1 tumors were undetectable by µCT (**Fig. 5A and B**) and only the 2 detectable tumors regrew upon termination of selpercatinib plus crizotinib treatment for 2 weeks (**Fig. 5B**), supporting complete elimination of 7 tumors by the up-front TKI combination. By contrast, all TR.2 tumors were detectable by µCT after 29 days of combination therapy and 6 of the 8 TR.2 tumors clearly progressed on treatment after another 20 days of treatment (**Fig. 5D**). The findings in **Figure 5** indicate that simultaneous inhibition of the Trim24-Ret oncogene and the HGF-MET bypass pathway yields superior therapeutic control in this murine lung cancer model relative to blocking the HGF-MET pathway at tumor progression.

**Figure 5.**
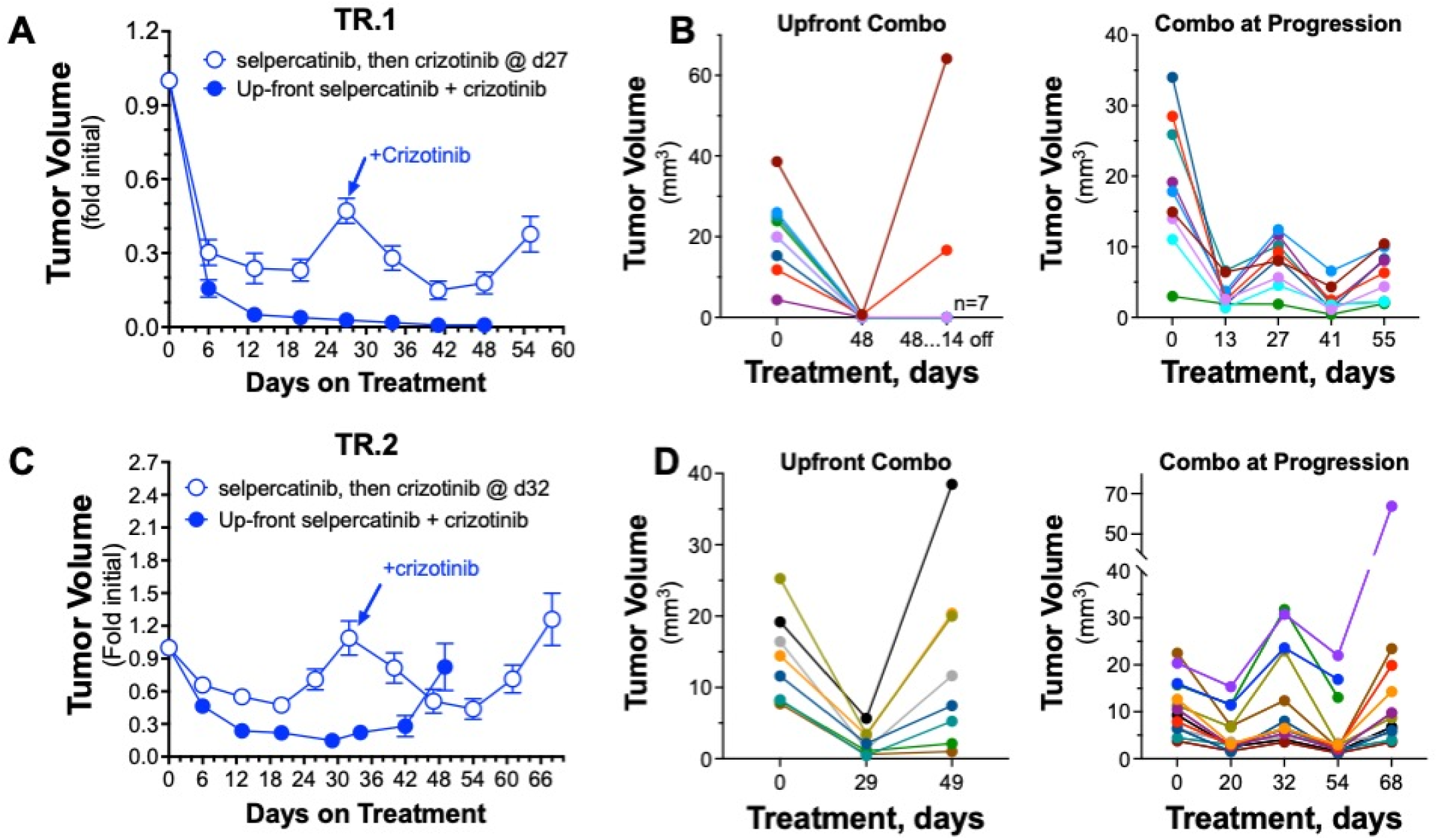
Therapeutic responses of orthotopic TR.1 and TR.2 tumors to selpercatinib and crizotinib combination treatment. TR.1 (**A, B**) or TR.2 (**C, D**) cells were injected into the left lungs of C57BL/6 mice as described in the Materials and Methods. After ∼10 days to allow tumor establishment, initial tumor volumes were measured by µCT and the mice were randomized into treatment groups (n=7-9 mice) that received selpercatinib alone (20 mg/kg) or combined selpercatinib and crizotinib (50 mg/kg). Mice were imaged weekly to monitor tumor volumes. When the TR.1 or TR.2 tumors in mice treated with selpercatinib alone had progressed (day 27 and 32, respectively), treatment was continued with combined selpercatinib and crizotinib. The data in **A** and **C** are presented as the mean and SEM of tumor sizes presented as fold of the initial pre-treatment tumor volumes. In **B** and **D**, the individual tumor volumes (mm^3^) in the upfront selpercatinib and crizotinib groups or the addition of crizotinib at progression on selpercatinib are shown at specific treatment dates from **A** and **C**.

## Discussion

The literature thoroughly documents the diverse mechanisms of acquired resistance to precision oncology agents including RET-targeting TKIs in lung cancer (7–9). Moreover, induction of MET signaling as a bypass resistance mechanism has been previously demonstrated in distinct subsets of RTK-driven lung cancers (12). Clinically, targeting MET in RTK-driven lung cancers progressing on the primary TKI with evidence of MET pathway activation yields responses, although usually partial with inevitable treatment failure (28–30). Likewise, when TR.1 and TR.2 tumors undergoing progression on single-agent selpercatinib treatment were switched to a combination of crizotinib and selpercatinib, significant but incomplete tumor shrinkage was observed, and the tumors again underwent progression (**Fig. 5**). Thus, the murine TR.1 and TR.2 orthotopic tumors accurately reflect the course of therapeutic responses frequently observed in lung cancer patients that acquire resistance to primary TKIs through MET pathway activity. It is important to note that despite equivalent *in vitro* control of TR.1 and TR.2 cells by upfront combination of selpercatinib and crizotinib (**Fig. 4**), different efficacies were observed upon treatment of orthotopic tumors derived from the cell lines (**Fig. 5**), suggesting that distinct paracrine interactions of the TR.1 and TR.2 tumor cells with the lung microenvironment contribute to the differing therapeutic responses. Our recent studies (15,16) with murine EGFRdel19 and EML4-ALK cell line models demonstrated important and variable contribution of host immunity to the overall TKI response measured *in vivo* and suggests that the different therapeutic efficacies of the upfront selpercatinib and crizotinib combination in TR.1 and TR.2 tumors may reflect differing recruitment of adaptive immune cells. The results reported herein and our recent publication therefore highlight the utility of murine lung cancer cell lines bearing relevant oncogene drivers and the orthotopic implantation model as a preclinical approach to molecularly define limitations of TKI monotherapy and test combination treatment strategies.

The incomplete clearing of lung tumor cells following treatment with oncogene-targeted agents yields residual disease or drug tolerant persisters and serves as a reservoir for selection and outgrowth of resistant cells. Many excellent reviews have clarified the key issues (31–33) and there is clear consensus within the field that combination therapies will be required to achieve the next significant advance in the treatment of oncogene-defined lung cancers (reviewed in (14,33)). The success of combination therapy in managing infectious diseases driven by tuberculosis and HIV provides important precedent (13) for combining agents that target the driving oncogene and emergent resistance mechanisms to achieve cures or long-term control in lung cancer. Like monotherapies in LUAD, serial treatment with single drugs targeting tuberculosis or HIV similarly failed and only succeeded when deployed as upfront cocktails. The key concept underlying the success of combination therapy for tuberculosis and HIV is prevention of the emergence of resistance (13). In this regard, the superior tumor control observed in the TR.1 and TR.2 orthotopic models with upfront selpercatinib and crizotinib relative to treating with crizotinib at progression (**Fig. 5**) suggests that this approach might prove fruitful in precision oncology-based therapies as well. Still, many challenges continue to hamper the development of highly effective combination therapies in oncogene-driven cancers. Defining the best drug combination to pursue in oncogene-defined cancers like LUAD remains a challenge. Also, implementation of upfront combination treatments in oncogene-driven lung cancers will require significant advances in biomarkers that flag a particular tumor for a specific drug combination.

The literature is replete with studies highlighting different combinations of oncogene-targeted agents and inhibitors of resistance, but how to prioritize them remains a hurdle to clinical translation. Identifying promising combinations of anticancer agents in preclinical studies invariably involves evaluation of drugs alone and in combination using high throughput *in vitro* platforms and short-term cell viability assays (3 to 7 days). In these experiments, priority is assigned to combinations that induce synergistic or synthetic lethal growth inhibitory activity. It is noteworthy that the *in vitro* combination of selpercatinib and the ERBB pathway inhibitor, afatinib yielded strong additive or synergistic reductions in TR.1 and TR.2 cell numbers when assayed at early time points (3-8 days of treatment) relative to combined selpercatinib and crizotinib at the same time points (see **Fig. 4**). Based on these data, a selpercatinib-afatinib combination might be prioritized for further study. However, the population doubling experiment performed unveiled the extended time course of the cellular response and demonstrated that *in vitro* acquired resistance to combined selpercatinib and afatinib occurred promptly in both TR.1 and TR.2 cells. By contrast, manifestation of the *in vitro* efficacy of combined selpercatinib and crizotinib required ∼14 days of treatment and reduced growth rates were subsequently sustained for the ∼40-day duration of the experiments (see **Fig. 4**). Moreover, upfront selpercatinib and crizotinib combination treatment of orthotopic TR.1 tumors yielded cures of the majority of the tumors relative to selpercatinib alone and significantly prolonged the time to progression in TR.2 tumors (**Fig. 5A** and **B**). We propose that the delayed onset of crizotinib action in combination with selpercatinib observed *in vitro* with TR.1 and TR.2 cells (**Fig. 4**) is due to the ability of the MET inhibitor to block function of a dominant and transcriptionally-induced MET signaling complex (**Suppl. Fig. S3** and **Fig. 3**) while the early action of afatinib may reflect rapid, non-transcriptional effects of EGFR-ERBB pathway-directed TKI resistance previously described in lung cancer cell lines bearing RTK fusion oncogenes (11). In light of this result, we wonder whether continued prioritization of drug combinations that induce synergistic cell killing in short-term *in vitro* assays may miss highly effective combinations of agents that prevent emergence and function of dominant bypass signaling pathways.

While the superiority of upfront selpercatinib and crizotinib combinations for treating orthotopic TR.1 and TR.2 tumors relative to addition of crizotinib at progression (**Fig. 5**) is compelling, rational deployment of this combination in lung cancer patients will require a measurement from clinical specimens that accurately identifies tumors destined to use HGF-MET pathway bypass signaling as a dominant resistance mechanism. MET gene copy number has been widely used as a marker of increased MET activity, but there is considerable debate concerning its utility as an accurate measure of MET biochemical activity (30,34,35). In fact, the literature indicates that MET gene copy number and levels of MET or pY-MET protein are poorly correlated in tumor specimens. Likewise, the levels of MET and pY-MET in selpercatinib-resistant TR.1 and TR.2 cell lines did not correlate well with MET inhibitor sensitivity. The observation that multiple MET and HGF interacting genes are increased in selpercatinib-resistant TR.1 and TR.2 cell lines supports the notion that increased activity of a MET signaling network through induction of HGF as well as specific co-receptors (PLXNB1, NRP1) and MET signaling adaptors like GAB1 may mediate increased MET activity. HGF expression was increased in some of the selpercatinib-resistant TR.1 cell lines (**Fig. 3A** and **C**), but not in selpercatinib-resistant TR.2 cell lines. In addition to autocrine HGF as a resistant mechanism, HGF is known to be expressed by distinct cell types within the tumor microenvironment and provide a paracrine mechanism mediating acquired MET-dependent resistance (36–38). A proximity ligation assay based on the interaction of MET and GRB2 demonstrated promise for assessing MET pathway activity in cell lines *in vitro* as well as patient specimens (35). Based on the likelihood that multiple interactors may increase overall MET network activity, evaluation of multiple interactors within the MET interaction network using proximity ligation assay-based approaches may identify a biomarker that accurately measures MET activity in clinical specimens.

## List of Abbreviations

RET: rearranged during transfection
MET: Mesenchymal Epithelial Transition
HGF: hepatocyte growth factor
ERBB: erythroblastic oncogene B
ALK: anaplastic lymphoma kinase
EGFR: epidermal growth factor receptor
RTK: receptor tyrosine kinase
µCT: microcomputed tomography
TKI: tyrosine kinase inhibitor
LUAD: lung adenocarcinoma

## Declarations

Ethics approval and consent to participate: All experiments were approved by the Institutional Animal Care and Use Committee at the University of Colorado Anschutz Medical Campus under an approved protocol (Precision Therapy in Lung Cancer #00860, approved February 10, 2025.

## Author Contributions

Concept and experimental design: TKH, LEH Developed methodology: TKH, RAN, ATL, LEH

Performed experiments and acquired data: TKH, TD, AA, SJ, SH, AN, LEH Analyzed and interpreted data: TKH, TD, AA, SJ, SH, LEH

Prepared figures and drafted manuscript: TKH, LEH

Edited and revised manuscripts: TKH, ATL, TD, AA, SJ, SH, AN, TP, RAN, LEH

## Availability of Data

The RNAseq data has been deposited in the GEO Datasets (GSE298990).

## Acknowledgements

The authors acknowledge the Genomics shared resource within the University of Colorado Cancer Center.

