## Supplemental Data for "Modeling acquired TKI resistance and effective combination therapeutic strategies in murine RET+ lung adenocarcinoma"

### Supplementary Table S1

#### Summary of IC50 values fromTKI Dose Response Experiments

| Cell Line | RET Selpercatinib | RET Pralsetinib | RET Embezotinib | MET Crizotinib | MET Capmatinib | ERBB family Sapitinib | ERBB family Afatinib | EGFR Gefitinib | FGFR AZD4547 | CSF1R Pexidartinib | MEK1/2 Trametinib | PTPN11 RMC-4550 | RAS RMC-7977 |
|---|---|---|---|---|---|---|---|---|---|---|---|---|---|
| | IC50, nM | | | | | | | | | | | | |
| TR.1 DMSO | 21 | 19 | 12 | >10 microM | >10 microM | >10 microM | >10 microM | >10 microM | >10 microM | ND | 58 | >10 microM | 20 |
| TR.1-1088 | 1365 | >10 microM | 398 | 405 | 4 | 9 | 87 | >10 microM | >10 microM | ND | 2 | 147 | 23 |
| TR.1-1090 | 1288 | 3284 | 554 | 1146 | 1 | 29 | 3 | >10 microM | >10 microM | ND | 0.7 | 103 | 34 |
| TR.1-1092 | >10 microM | >10 microM | >10 microM | 168 | 5 | 1022 | 727 | >10 microM | >10 microM | ND | 2 | 55 | 36 |
| TR.1-1094 | >10 microM | >10 microM | >10 microM | 191 | 7 | 6 | 34 | >10 microM | >10 microM | ND | 2 | 57 | 27 |
| TR.2 DMSO | 16 | 15 | 19 | >10 microM | >10 microM | >10 microM | 774 | ND | ND | >10 microM | 5 | >10 microM | 6 |
| TR.2-A0827 | 1340 | 291 | 460 | 532 | <1 nM | 9 | 4 | ND | ND | >10 microM | 0.3 | 13 | 8 |
| TR.2-A0828 | 1099 | 297 | 342 | 520 | <1 nM | 5 | 3 | ND | ND | >10 microM | 0.4 | 16 | 10 |
| TR.2-A0831 | 2596 | 1281 | 599 | 273 | <1 nM | 6 | 4 | ND | ND | >10 microM | 0.4 | 44 | 9 |
| TR.2-A0832 | 6718 | 535 | 1309 | 502 | <1 nM | 9 | 5 | ND | ND | >10 microM | 0.4 | 48 | 6 |

ND, not determined

**Supplementary Table S1**

**Supplementary Table S1. $IC_{50}$ values for dose-response data in Figures 1 and 2.** The $IC_{50}$ values associated with the dose-response experiments presented in Figures 1 and 2 were calculated with the Prism 10 software program using nonlinear curve-fitting and the log(inhibitor) vs. normalized response model and robust regression.

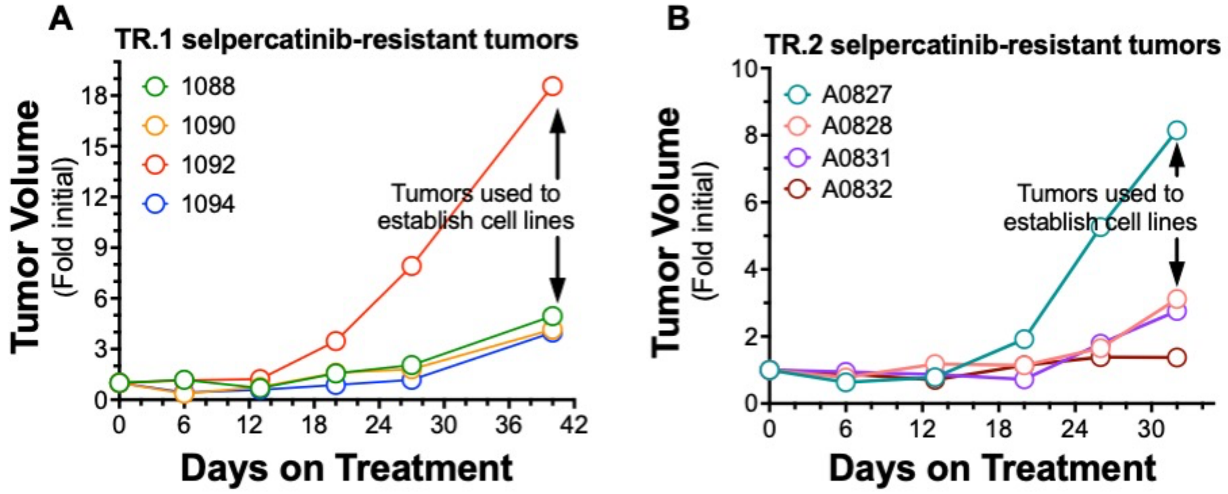


Supplementary Figure S1

**Supplementary Figure S1. Orthotopic TR.1 and TR.2 cell-derived tumors progress on continuous selpercatinib treatment.** TR.1 (**A**) and TR.2 (**B**) cells were inoculated into the left lungs of C57BL/6 mice as described in the Materials and Methods. After 10 days, the mice were submitted to μCT imaging and pretreatment tumor volumes were obtained. The mice were treated with selpercatinib (daily gavage for 5 days and 2 days off) and tumor volumes were measured with μCT weekly. Individual tumor volumes as fold of the initial volumes are presented. The mice were euthanized after 40 days (TR.1) or 32 days (TR.2) of treatment and the excised tumors were used to propagate TKI-resistant cell cultures in growth medium containing 200 nM selpercatinib.

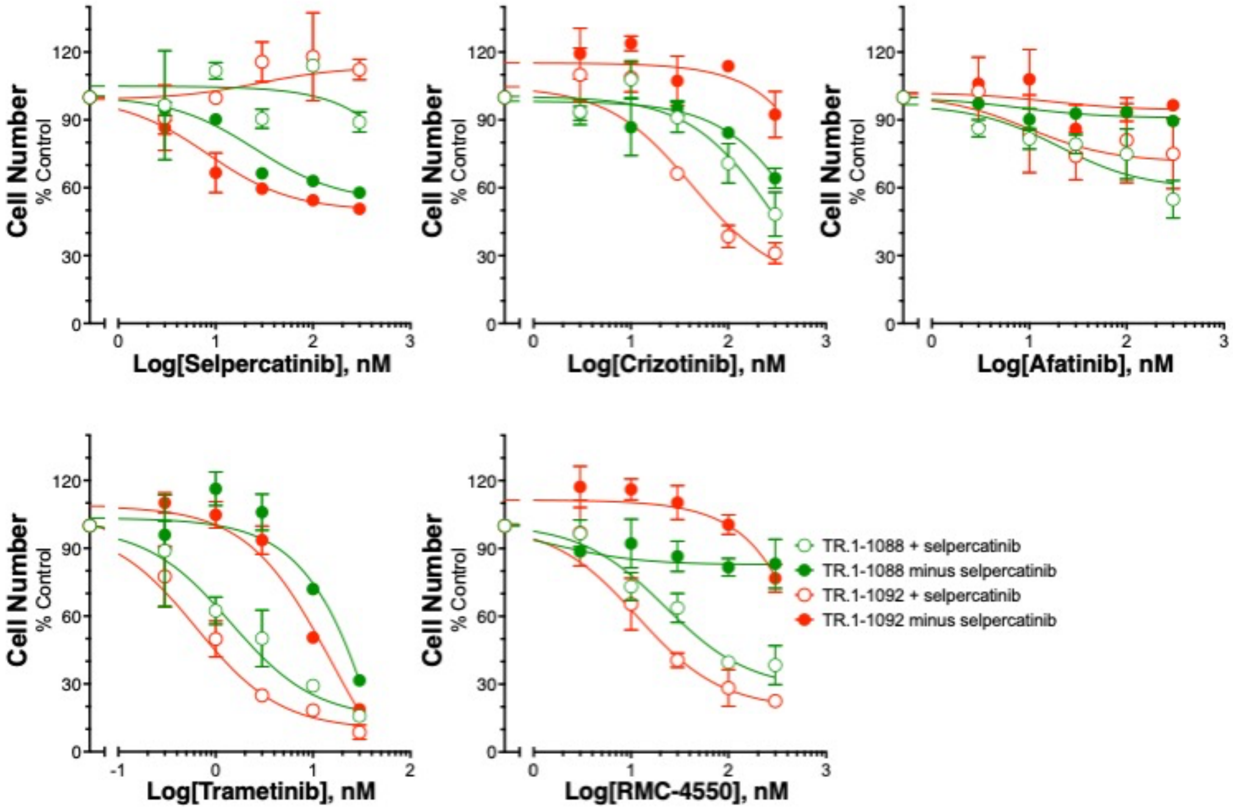


nentary Figure S2

**Supplementary Figure S2. Partial reversibility of selpercatinib-resistant drug sensitivities.** TR.1-1088 and -1092 selpercatinib-resistant cell lines were cultured in the absence of selpercatinib for 14 days and submitted to 96-well dose-response experiments with the indicated agents. TR.1-1088 and -1092 cells cultured for 14 days with 200 nM selpercatinib were tested as controls. After 7 days of incubation, relative cell numbers were assayed with CyQUANT reagent as described in the Materials and Methods.

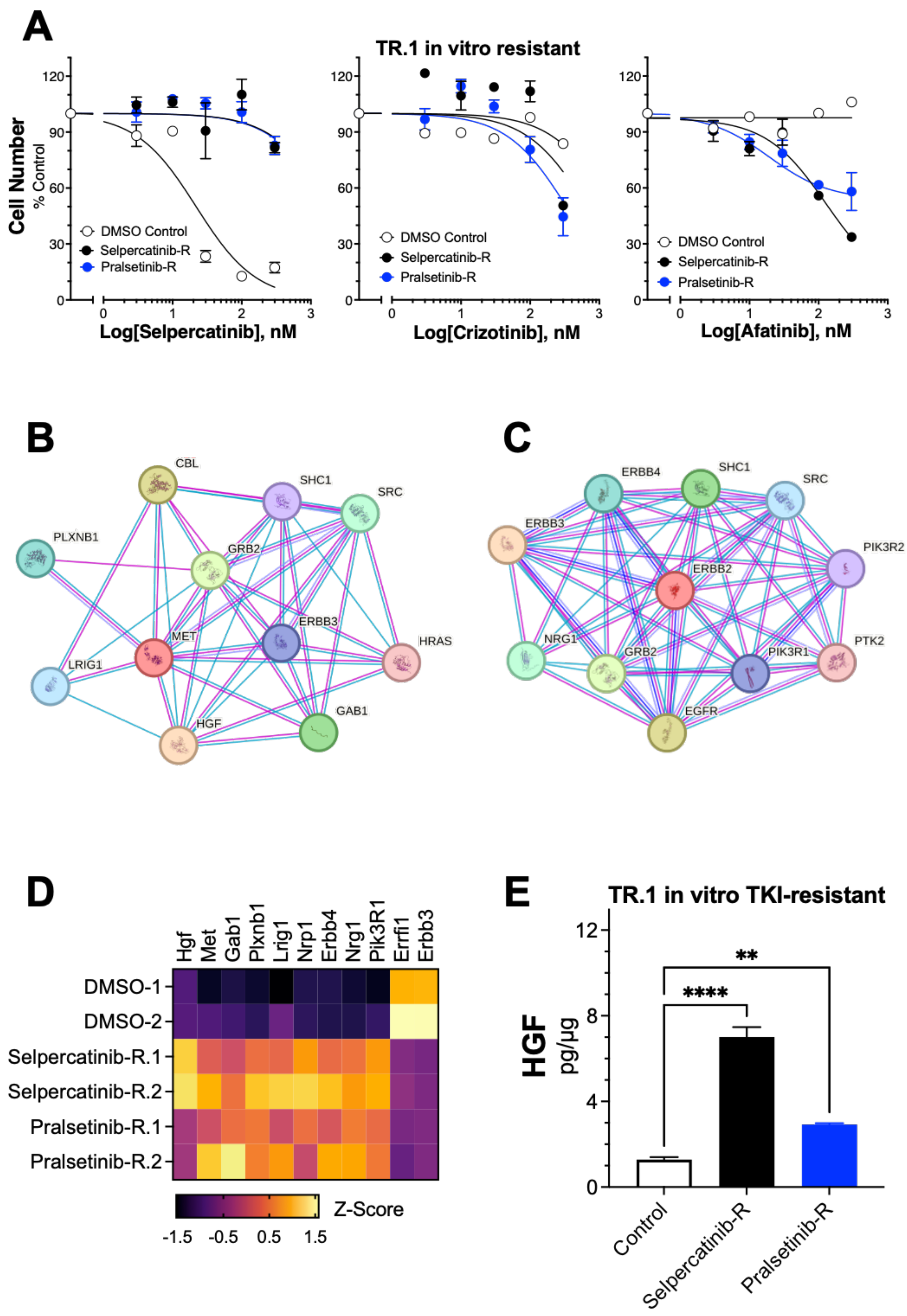


**Supplementary Figure S3**

**Supplementary Figure S3. Analysis of TR.1 cells made resistant to selpercatinib and pralsetinib *in vitro* reveals evidence for induction of MET and ERBB4 interaction networks.** TR.1 cells were made resistant to selpercatinib and pralsetinib using *in vitro* culture methods and dose escalation of TKI concentrations. **A**, The TKI-resistant cell cultures as well as TR.1 cells cultured in 0.1% DMSO as a passage control were tested for sensitivity to selpercatinib, crizotinib and afatinib in a 96-well format. After 7 days of incubation, cell numbers were measured with CyQUANT reagent and the data are presented as percent of DMSO-treated cells. The predicted MET (**B**) and ERBB4 (**C**) interaction networks were determined using the STRING database (string-db.org/). The queries were limited to experiments, databases and co-occurrence active interaction sources, but not text mining. **D**, The RNAseq data were queried for mRNA levels of the indicated MET and ERBB4 interacting genes, normalized to Z-scores and presented in a heatmap format. **E**, Conditioned media from the TR.1 cell lines made TKI resistant *in vitro* were submitted to ELISA for HGF. The values were normalized to cellular protein and presented as the means and SEM of three independent experiments.

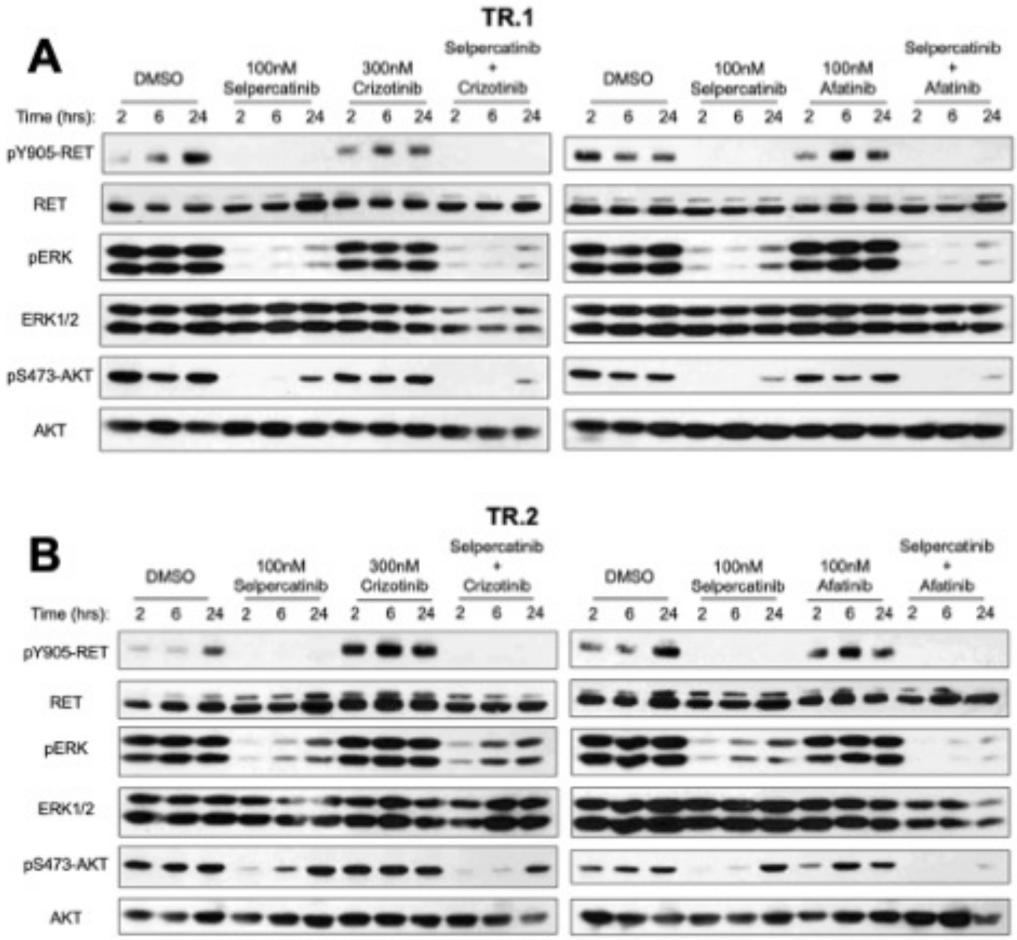


**Supplementary Figure S4**

**Supplementary Figure S4. Immunoblot analysis of ERK MAPK and AKT in TR.1 and TR.2 cells treated with selpercatinib alone and in combination with crizotinib or afatinib.** TR.1 and TR.2 cells in full growth medium were treated for 2, 6 and 24 hours with 100 nM selpercatinib, 300 nM crizotinib or 100 nM afatinib alone or in combination. Cell-free extracts were prepared as described in the Materials and Methods and submitted to SDS-PAGE. The gels were electrophoretically transferred and probed with the indicated antibodies.
